# Changes in motor unit conduction velocity after unilateral lower limb suspension and active recovery correlate with muscle ion channel gene expression

**DOI:** 10.1101/2025.02.26.640329

**Authors:** Giacomo Valli, Fabio Sarto, Francesco Negro, Elena Monti, Giuseppe Sirago, Matteo Paganini, Sandra Zampieri, Martino V. Franchi, Andrea Casolo, Julián Candia, Luigi Ferrucci, Marco V. Narici, Giuseppe De Vito

## Abstract

The effects of muscle disuse on the propagation of action potentials along muscle units, a key process for effective muscle activation and force production, remain poorly understood. This study aimed to investigate changes in action potential propagation and to identify biological factors influencing these changes following unilateral lower limb suspension (ULLS) and active recovery (AR). Eleven young male participants underwent 10 days of ULLS followed by 21 days of AR based on resistance exercise. Maximal force of the knee extensor muscles (MVC), High-Density surface EMG recordings and muscle biopsies of the *vastus lateralis* muscle were collected before ULLS, after ULLS and after AR. EMG recordings collected during submaximal isometric contractions were decomposed to estimate single motor unit conduction velocity (MU CV). Muscle biopsies were used to measure muscle fibre diameters via histochemical analysis and ion channel transcriptomic profiles via mRNA-sequencing. MVC decreased after ULLS by 29% and fully recovered after AR. MU CV decreased after ULLS and fully recovered, up to exceeding baseline values after AR. Muscle fibre diameters did not change across the interventions and showed no correlation with MU CV. Conversely, a feature importance analysis revealed that mRNA expression levels of specific ion channel genes, particularly those involved in K^+^ transport, correlate with MU CV at baseline and across the interventions. This study highlights the crucial role of K^+^ ion channels in influencing MU CV in humans, offering new insights into MU CV modulation and the mechanisms of muscle force changes after disuse and active recovery.

**Key points:**

- Muscle disuse, such as in unilateral lower limb suspension, leads to a decrease in motor unit conduction velocity (MU CV), a critical factor for muscle activation and force production.
- Active recovery through resistance exercise results in the full recovery of MU CV, even exceeding baseline levels.
- Muscle fibre diameters do not change significantly after limb suspension or active recovery and show no correlation with MU CV.
- Conversely, ion channel mRNA expression, particularly of those related to K^+^ transport, correlates with MU CV and its changes following disuse and recovery.
- These findings highlight K^+^ ion channels as a key factor in regulating MU CV in humans and provide new molecular determinants of the changes in muscle force after disuse and recovery.

## 1. Introduction

Experimental models of muscle disuse are widely adopted to investigate the deleterious consequences of micro-gravity or hospitalisation-induced inactivity (Qaisar *et al*., 2020) and to study their underlying mechanisms and potential countermeasures (Michel *et al*., 2024; Franchi *et al*., 2025). Disuse commonly leads to a reduction in both muscle mass and function, with the decline in muscle function significantly exceeding the loss of muscle mass (Campbell *et al*., 2019; Monti *et al*., 2021). This disparity suggests a pivotal role of neuromuscular factors in driving functional impairments (Piasecki, 2024).

Despite its critical role in controlling force production, neuromuscular function has received relatively little attention in disuse research, particularly regarding motor unit (MU) properties. In fact, until recent years, only a few studies investigated the changes in MU properties induced by muscle disuse. These studies observed a reduced MU discharge rate in small hand muscles after hand cast immobilisation (Duchateau & Hainaut, 1990; Seki *et al*., 2001, 2007) and a reduced speed of action potential propagation along muscle fibres in large leg muscles after bed rest (Cescon & Gazzoni, 2010). Although these previous studies built the foundation for understanding the impact of muscle disuse on neuromuscular function, several important questions remain unanswered, largely due to the limitations of the techniques available at the time.

Thanks to technological advancements in neuromuscular assessment techniques (Farina *et al*., 2016), recent studies employed state-of-the-art Intramuscular and High-Density surface Electromyography (HDsEMG) to gain new insights into MU behaviour and neuromuscular adaptations during muscle disuse and recovery (Sarto *et al*., 2022*b*; Piasecki, 2024).

For instance, a study from our group (Valli *et al*., 2024*b*) shed light on the longstanding question of whether the reduction in MU discharge rate is preferentially observed in lower-threshold MUs, as suggested by (Duchateau & Hainaut, 1990), or affects all MUs uniformly (Seki *et al*., 2001). Our findings based on a large sample of MUs showed that after 10 days of unilateral lower limb suspension (ULLS), the reduction in discharge rate is indeed specific to lower-threshold MUs, suggesting that this selective decrement of neural drive may lead the shift in fibre type phenotype (from slow to fast) that is commonly observed after longer periods of disuse (Bodine, 2013).

Other studies focused instead on understanding the mechanisms behind the disuse-induced neuromuscular adaptations. Focusing on MU central properties, it was shown that neuromodulation decreases following ULLS, which may partially explain the diminished neural drive to the muscle (Martino *et al*., 2024). On the other hand, studies performed with intramuscular EMG have shown that the neuromuscular junction is functionally resilient to 10-15 days of muscle unloading (Sarto *et al*., 2022*a*) or immobilisation (Inns *et al*., 2022) and, therefore, it remains capable of reliably conveying the neural discharge to the muscle fibres within this time-frame.

In summary, the neuromuscular consequences of disuse have been characterised at the spinal, motoneuronal and neuromuscular junction level. However, modifications at the final component of the neuromuscular system, the muscle unit (Heckman & Enoka, 2004), remain to be fully elucidated.

The aim of this study is to clarify the modifications occurring at the muscular component of the MU following 10 days of ULLS and subsequent active recovery (AR), and to identify possible factors involved in these adaptations. To achieve this, we used HDsEMG recordings to study the conduction velocity (CV) of single MU action potentials (MUAPs), and muscle biopsies to investigate muscle fibre diameters and ion channel mRNA expression. We focused on MU CV, which represents the speed at which an action potential travels along the muscle fibres of a single MU, as it is a crucial marker of MU contractile properties (Andreassen & Arendt-Nielsen, 1987). From a biophysical perspective, it has been observed that CV of electrically evoked action potentials correlates with *vastus lateralis* muscle fibre size (Methenitis *et al*., 2016), although the molecular determinants of MU CV in humans remain poorly understood. To address this, we combined the analysis of muscle fibre diameters and ion channel mRNA expression, as these channels play a crucial role in the generation and propagation of action potentials and may significantly influence MU CV and its changes (Jurkat-Rott & Lehmann-Horn, 2004).

We hypothesised that MU CV would decrease after ULLS and recover after AR. We further posited that these changes in MU CV could be explained by alterations in muscle fibre diameters and/or ion channel mRNA expression.

## 2. Methods

This study is part of a larger investigation focused on identifying early biomarkers of neuromuscular degeneration following short-term unloading. For a detailed explanation of the unloading and active recovery models and biological data extraction, the reader can refer to (Sarto *et al*., 2022*a*), and for a detailed explanation of the HDsEMG data collection to (Valli *et al*., 2024*b*).

### 2.1. Ethical approval

This study was approved by the Ethics Committee of the Department of Biomedical Sciences of the University of Padova (Italy) with reference number HEC-DSB/01-18. The study was conducted in accordance with the standards set by the latest revision of the Declaration of Helsinki. Participants were informed about all the experimental procedures through an interview and information sheets. Their eligibility was determined following a thorough review of their medical history. Volunteers were enrolled in the study after signing a written consent form, with the option to withdraw at any time.

### 2.2. Participants and experimental protocol

Twelve healthy, recreationally active young male adults (age: 22.1 (2.9) years; height: 1.78 (0.03) m; body mass: 72.1 (7.1) kg) volunteered for this study. To minimize the risk of deep venous thrombosis associated with ULLS (Bleeker *et al*., 2004), only male participants were included, as among young individuals the absolute risk of first venous thrombosis is higher in females (Roach *et al*., 2014). The inclusion criteria were: age between 18 and 35 years, a body mass index of 20 to 28 kg/m², and engagement in recreational physical activities 1 to 3 times per week (self-reported). Exclusion criteria included a sedentary lifestyle, participation in recreational physical activities more than 3 times per week, smoking, a history of deep venous thrombosis, and any other conditions that might prevent safe participation in the study.

Data collection was performed at baseline (day 0 of limb suspension, LS0), after 10 days of ULLS (LS10), and following 21 days of active recovery (AR21) (Fig. 1A). All the participants visited the laboratory prior to the LS0 data collection to familiarise with the ULLS procedures and with the isometric muscle contractions. At LS10, participants were tested immediately after ending limb suspension and were only allowed to warm up before the MVC test. At AR21, tests were conducted approximately 72 hours after the final exercise session to avoid potential muscle fatigue.

**Figure 1:**
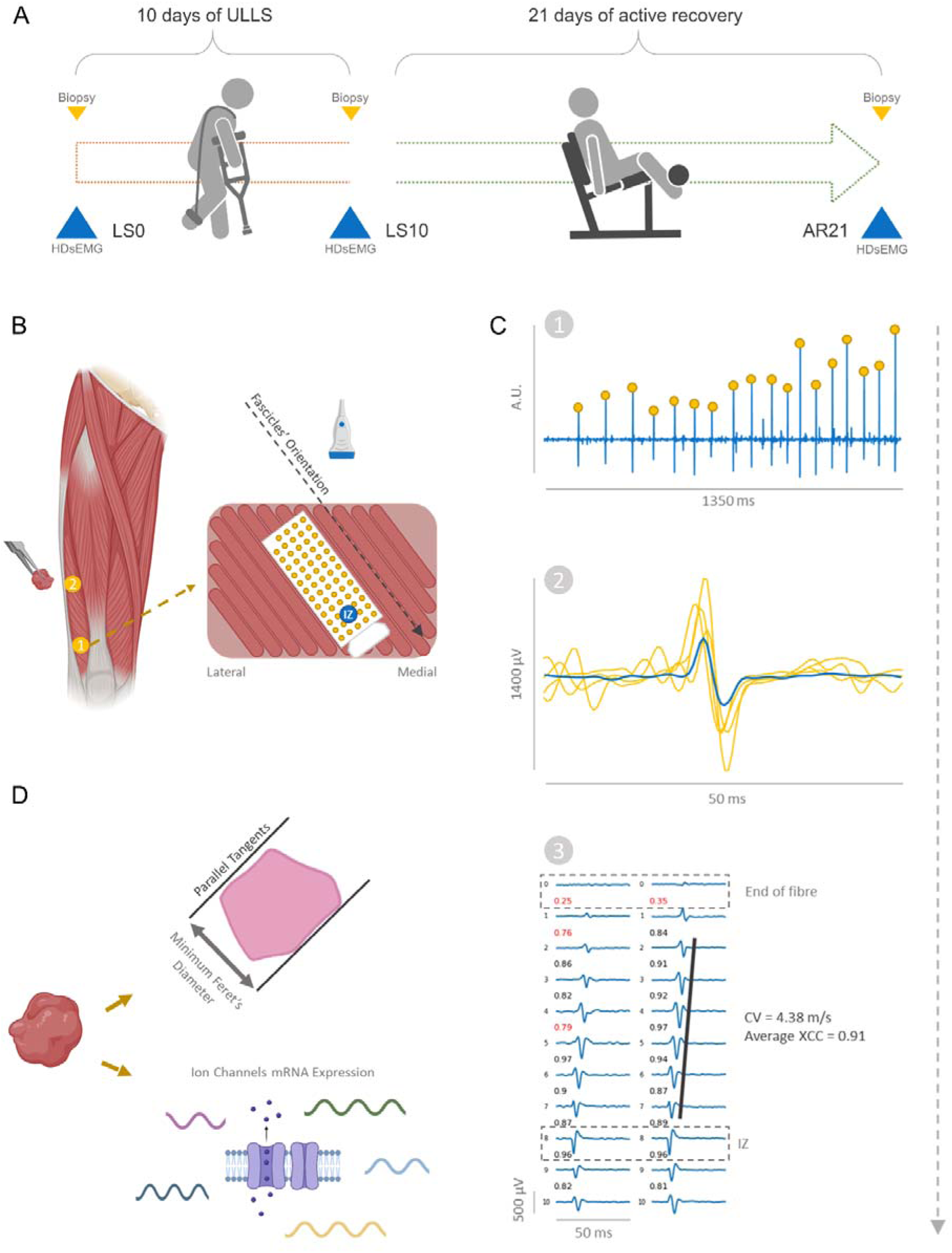
Schematic representation of the study design (A) and procedures of data collection and analysis (B-D). Data was collected at baseline (day 0 of limb suspension, LS0), after 10 days of ULLS (LS10) and after 21 days of active recovery (AR21) (A). Panel B shows the expected location of the used innervation zones (IZ) in the vastus lateralis muscle. The central zone (around the IZ number 2), was used for the biopsy collection while the distal zone (around the IZ number 1) was used for the High-Density surface Electromyography (HDsEMG) recordings. Panel B also shows the orientation of the recording grid with respect to the muscle fascicles and the IZ. The procedure used for the extraction of motor unit action potentials (MUAPs) and for MU conduction velocity (CV) estimation is depicted in panel C. Specifically, the time of firing of each MU (C1) was used to trigger and average the MUAPs in each channel of the grid (C2) and the best propagating channels were selected for MU CV estimation via a maximum likelihood algorithm (C3). Muscle biopsies were used for the estimation of the minimum Feret’s diameter and for the measurement of the Ion channel gene expression via bulk RNA sequencing. XCC, cross-correlation coefficient, is the measure of similarity between the MUs action potential shape.

Participants were instructed to avoid intense exercise, coffee, and alcohol for 24 hours before any data collection. All tests were performed at the same time of day for each participant.

### 2.3. Intervention

#### 2.3.1. Unilateral lower limb suspension

The dominant lower limb (the kicking leg, which was the right leg for all participants) was suspended in a slightly flexed position (15 to 20 degrees of knee flexion). To ensure the suspended limb did not touch the ground while moving, the left lower limb was fitted with an elevated shoe (50 mm sole) (Berg *et al*., 1991). Throughout the ULLS period, participants walked exclusively using crutches and avoided any weight-bearing on the unloaded limb. Participants were instructed to avoid any active contraction of the suspended limb’s muscles. However, they were allowed to perform passive, non-weight-bearing ankle exercises (Bleeker *et al*., 2004) to promote blood circulation and reduce the risk of deep venous thrombosis. Participants were recommended to wear elastic compression socks on the suspended lower limb during ULLS. Participant’s compliance was evaluated through daily calls and messages.

#### 2.3.2. Active recovery

The AR phase began approximately 72 hours after the suspension period and lasted 21 days. During the AR phase, participants performed unilateral resistance exercises three times per week, with at least 24 hours of rest between sessions. Each session included 3 sets of 10 repetitions of leg press and leg extension exercises at 70% of the estimated 1 repetition maximum (1RM), following a warm-up at 30% 1RM. Exercises were performed from full knee extension to ∼90° flexion with a time under tension of ∼2 seconds both in the concentric and eccentric phases. A 2-minute rest was observed between sets.

1RM was estimated from the heaviest load that participants could lift for 4-6 repetitions during the first session (Brzycki, 1993), and was re-evaluated weekly to adjust the load. This approach was chosen due to participants’ lack of prior resistance training and the recent unloading of their lower limb.

### 2.4. Measurements

#### 2.4.1. High-density surface electromyography grid placement

HDsEMG signals were recorded from the distal portion of the *vastus lateralis* muscle (Fig. 1B) using an adhesive grid of 64 electrodes (5 columns and 13 rows with 8-mm interelectrode distance (GR08MM1305, OT Bioelettronica, Torino, Italy) filled with conductive cream (Ac cream, OT Bioelettronica, Torino, Italy).

A meticulous 2-steps approach has been adopted to ensure the precise alignment of the grid’s columns with respect to the muscle fibres and to maintain a standardised position of the grid with respect to the muscle innervation zone, as these are fundamental prerequisites for reliable MU CV estimation (Valli *et al*., 2024*a*). First, the innervation zone was detected as the location in which the largest muscle twitch was evoked in response to low-intensity percutaneous electrical stimulation using a pen electrode (Botter *et al*., 2011). Since *vastus lateralis* presents different innervation zones, we focused on the most distal one which is generally located between 35% and 20% of femur length and provides better MU decomposition (Botter *et al*., 2011) (Fig 1.B). Stimulations were induced using an electrical stimulator (DS7AH, Digitimer Ltd, Welwyn Garden, Hertfordshire, UK) with an electrical current set at 16 mA (400V, pulse width: 50 µs). Once the innervation zone was identified, the current was reduced to 8-10 mA to narrow the identified location. Second, muscle fascicles orientation over the innervation zone was detected using B-mode ultrasound recordings (Mylab70, Esaote, Genoa, Italy) (Hug *et al*., 2021) and it was marked on the skin with a permanent marker.

Subsequently, the grid was placed following the muscle fascicles orientation and above the innervation zone (Botter *et al*., 2011; Barbero *et al*., 2012). Specifically, the innervation zone corresponded with the central electrodes of the last two-three rows of the grid. Please refer to Fig. 1B for a visual representation.

The ultrasound system was used also to accurately detect muscle borders and avoid the placement of the grid across adjacent muscles.

Before positioning the grid, the skin was carefully shaved, cleaned with 70% ethanol, and then treated with abrasive-conductive paste (Spes medica, Salerno, Italy). Two reference electrodes were placed on the malleolus and patella bones.

After the recordings, the grid border was marked with a permanent marker on the skin. The operator emphasised these markings at each participant meeting to ensure that the grid’s exact placement could be reproduced in the subsequent data collection points (Casolo *et al*., 2020).

#### 2.4.2. High-density surface electromyography recordings

Maximum voluntary contraction (MVC) of the knee extensor muscles was assessed at 90° knee angle using a custom-made knee dynamometer fitted with a load cell (RS 206–0290) which was attached above the ankle through straps. The participant’s back was supported in an upright position, with a resulting hip angle of 90°. The hip was stabilised to the table with adjustable straps to limit compensation (Monti *et al*., 2021). Participants were instructed to perform the task during the familiarisation session and, after a standardised warm-up, were asked to “push as hard as possible” by extending the dominant leg against the load cell, and then to maintain the contraction for 3-4 seconds. Loud verbal encouragement and visual feedback were provided. The test was repeated three times with 60 seconds of rest and the contraction with the maximum value was used to determine the target forces used for HDsEMG recordings.

The HDsEMG signal was recorded during submaximal isometric trapezoidal contractions performed at 10, 25 and 50% MVC. Each contraction consisted of a ramp-up phase to the submaximal target force level which was then maintained during a steady-state phase, and then a ramp-down phase back to baseline. All the contractions had a total duration of 30 seconds. The ramp-up and ramp-down phases were performed with a linear force increase/decrease set at 5% MVC per second (Del Vecchio *et al*., 2019) and the duration of the steady-state phase was adjusted accordingly (i.e., at 50% MVC the ramps lasted 10 seconds each and the steady-state 10 seconds, which adds up to 30 seconds of total contraction). Each intensity level of the trapezoidal contractions was repeated twice, in random order, with 60 seconds of rest in between. Participants received real-time visual feedback of the force produced and were instructed to match it as precisely as possible during the familiarisation session.

The EMG and force signals were sampled at 2048 Hz with the EMG-Quattrocento (OT Bioelettronica, Torino, Italy). The EMG signal was recorded in monopolar configuration, amplified (×150) and band-pass filtered (10–500 Hz) at the source. Force was recorded synchronously with the EMG signal and the offset was removed before starting the recording.

### 2.5. Motor unit detection and conduction velocity estimation

The force signal was low-pass filtered using a fourth-order, zero-lag Butterworth filter with a 15 Hz cut-off. The HDsEMG signal was band-pass filtered between 20 and 500 Hz with a second-order, zero-lag Butterworth filter and decomposed to obtain the discharge pattern of individual MUs with convolutive blind source separation (Fig. 1C-1) (Negro *et al*., 2016). The majority of the MUs included in this study are the same as in our previous publication (Valli *et al*., 2024*b*). However, the decomposition of some contractions with a limited number of detected MUs was repeated with the aim to maximise the number of MUs suitable for CV analysis. Specifically, particular attention was dedicated to the removal of noisy channels and the decomposition was run until the residual activity index reached zero, rather than up to a fixed number of iterations. This additional decomposition allowed also for the inclusion of one more participant in the analyses compared to our previous publication (Valli *et al*., 2024*b*). After decomposition, the discharge pattern was inspected and manually edited by an experienced operator following recent guidelines (Del Vecchio *et al*., 2020; Martinez-Valdes *et al*., 2023) and only MUs with a pulse to noise ratio (PNR) ≥ 28 were maintained for further analyses (Holobar *et al*., 2014).

MUs detected during the two contractions performed at the same intensity level during the same data collection point were pooled and analysed together after the removal of duplicated MUs as previously explained (Valli *et al*., 2024*b*). Briefly, the MUs detected in both contractions where matched via comparison of their MUAPs representation and those with a cross-correlation coefficient (XCC) above 0.9 where classified as duplicates (Maathuis *et al*., 2008; Martinez-Valdes *et al*., 2017). Of the 2 duplicates, the MU with the lowest PNR was removed. This approach allowed us to obtain a broader and more representative number of MUs to include in the following analysis, an important aspect especially considering the strict criteria used for the selection of MUs suitable for CV analysis.

For MU CV estimation, the MUAPs representation across the grid’s channels was obtained from spike-triggered averaging of the double-differential derivation of the EMG signal along the direction of the muscle fibres (Fig. 1C-2) (Casolo *et al*., 2020). This spatial filtering is important to enhance the representation of MUAPs propagation as it decreases the presence of non-propagating components and attenuates the end-of-fibre effect (Gallina *et al*., 2022). The spike-triggered averaging was performed using the first 50 MU discharges (Martinez-Valdes *et al*., 2018) and with a 50 milliseconds window (Valli *et al*., 2024*c*, 2024*a*).

The channels used for MU CV estimation were manually selected by an experienced operator using the following inclusion criteria: i) a clear propagation of the action potential, ii) an XCC between adjacent channels above 0.8 and iii) no innervation zone or end of fibre effect (Fig. 1C-3). If multiple columns presented similarly suitable MUAPs, the selection prioritised the column with the highest average XCC between adjacent channels (Valli *et al*., 2024*a*). On the selected channels, MU CV was calculated using the maximum likelihood estimation of delay previously proposed by (Farina *et al*., 2000, 2002). Although this maximum likelihood algorithm can theoretically work with a minimum of two signals, we always selected the greatest possible number of channels in order to maximise the accuracy of the estimation.

For the scope of this study, we only investigated the total pool of decomposed MUs without applying any longitudinal tracking procedure (Martinez-Valdes *et al*., 2017). This was preferred because MU tracking across 3 timepoints, with ULLS and AR between them, greatly reduces the number of MUs suitable for analysis (to less than 20% of the total pool) (Valli *et al*., 2024*b*) which would be further reduced after applying the inclusion criteria required for accurate MU CV estimation. Therefore, channels selection for MU CV estimation was performed independently for each MU at each data collection point, with the operator blind to the condition.

For correlation analyses, the participants’ mean MU CV values have been obtained by averaging the mean MU CV at the three contraction intensities (i.e., mean MU CV = (mean MU CV at 10% MVC + mean MU CV at 25% MVC + mean MU CV at 50% MVC) / 3)). This method was chosen over a global average of all MU CV values, as it avoids bias from differences in the number of MUs detected at each contraction intensity (Casolo *et al*., 2023). Throughout this manuscript, “mean MU CV” will refer to this calculated average. Mean CV was used for all the correlation analyses except for the correlation between MU RT and CV, as explained in the “Statistical analysis” section.

For the MUs were CV could be estimated, the relative MU recruitment threshold (RT) was also calculated (i.e., the relative (% MVC) force level corresponding to the first discharge time).

MU decomposition was performed with custom Matlab scripts (R2023a; The Mathworks Inc., Natick, MA), while all the analyses were performed with the *openhdemg* V0.1.0 library (Valli *et al*., 2024*a*) in Python (V3.11.6, Python Software Foundation, USA).

### 2.6. Muscle biopsy collection and muscle fibre diameters estimation

From each participant, a muscle biopsy of approximately 150 mg was collected from the *vastus lateralis* muscle using a Weil–Blakesley conchotome (Gebrüder Zepf Medizintechnik GmbH & Co. KG, Dürbheim, Germany). The three biopsies were performed at 2 cm from the central innervation zone and with a distance between each-others of about 2-3 cm to avoid effects of pre-sampling (Sarto *et al*., 2022*a*). With this setup, the bandage of the biopsy (and consecutive swelling) would not affect HDsEMG recordings, which are instead performed on the distal innervation zone. Furthermore, given that the longitudinal position of the biopsy site does not affect biological properties (including muscle fibre diameters) in the *vastus lateralis*, (Horwath *et al*., 2021) the use of different sites for the collection of biopsies and HDsEMG recordings should not introduce any bias in the relation between biological and electrophysiological parameters.

Within 3 to 5 minutes from data collection, a portion (∼30 mg) of the muscle was cleaned by connective and adipose tissue, embedded in an oriented way in optimal cutting temperature (OCT) compound, frozen in isopentane, and stored at −80°C for histochemical analysis. Cryosections were made using a manual cryostat (Leica CM1850; Leica Microsystems, Wetzlar, Germany), producing sections that were 10 μm thick. To evaluate muscle fibre diameter depending on fibres type, muscle sections were stained for myofibrillar ATPases with pre-incubation at pH 4.35 on serial cross-sections (Carraro *et al*., 1985) resulting in slow-twitch muscle fibres (with slower ATPase activity) appearing dark, while fast-twitch fibres (with faster ATPase activity) were lightly stained.

Since the estimation of muscle fibres diameter can be affected by the orientation of the sectioning angle or by kinked muscle fibres (Dubowitz *et al*., 2013), we opted to estimate fibre diameters via measuring the minimal Feret’s diameter, defined as the minimum distance of parallel tangents at opposing borders of the muscle fibre. Indeed, this approach can minimise the bias introduced by the variable cutting of the sections (Briguet *et al*., 2004). The minimum Feret diameter of each muscle fibre was manually measured across all visible fibres (approximately 200–400 per section) using the ImageJ software v1.52v (Schindelin *et al*., 2015).

### 2.7. Ion channel mRNA expression

Full description of mRNA sequencing data extraction starting from the biopsies can be found in (Sarto *et al*., 2022*a*). For the scope of this study, mRNA sequencing raw counts have been normalised using the R package DESeq2 V4.4.1 (Love *et al*., 2014) called from Python via the rpy2 package V3.5.16 and associated with the corresponding metadata (i.e., participant and data collection point). Of all the available genes, we extracted those present in skeletal muscle ion channels based on (Jurkat-Rott & Lehmann-Horn, 2004), which resulted in 35 genes suitable for the following analyses.

### 2.8. Statistical analysis

For MVC, muscle fibre diameters (slow fibre, fast fibre and total fibre diameters) and mRNA expression of selected ion channel genes (*KCNJ2-AS1*, *KCNN2*, *KCNN3*, *SCN4A*), variables normality of the distribution was assessed through the Shapiro–Wilk test. Since the normality assumption was satisfied, a one-way repeated measure analysis of variance (ANOVA) was used. For all ANOVAs, sphericity was tested with Mauchly’s test and if the assumption of sphericity was violated, the correction of Greenhouse-Geisser was applied. If the ANOVA was significant, the Post Hoc pairwise T-tests with Holm correction were used to determine whether differences among the time points were present. Partial eta squared (□p^2^) was also estimated to evaluate the effect size for ANOVAs.

MU CV was analysed using a linear mixed-effects model, as multiple MUs were detected from each participant (Yu *et al*., 2022). The normality of the model residuals was assessed through visual inspection of the Q-Q plot and histogram. Given that normality of the residuals was confirmed, the linear model was applied with “Time” and “Intensity” as factors and the “Participant” as a cluster variable. Post Hoc comparisons were performed with Holm correction.

A linear mixed-effects model was also applied to investigate the relationship between MU RT and CV and to determine whether this relationship varied across the different data collection points. Specifically, the latter was assessed by comparing changes in the slopes and intercepts of the regression lines for mean MU RT and CV values at each data collection point. For this analysis, we used the mean RT and CV values at each contraction intensity (i.e., three values per participant, one each at 10%, 25%, and 50% MVC) instead of the previously described overall mean RT and CV. This choice helped minimising overfitting in the correlation analysis and to capture a broader range of values. Additionally, this approach yields similar correlation values to the mean within-participant correlation method used by (Casolo *et al*., 2023) (data not shown) while allowing for a simpler model to compare slopes and intercepts across data collection points.

Repeated measures correlation (Bakdash & Marusich, 2017) was used to determine whether changes in mean MU CV across the three data collection points were associated with changes in MVC, total muscle fibre diameters and ion channel mRNA expression. For this analysis, average values for each participant MU CV were used as a representation of the clustered values in order to reduce the variability of the model as previously suggested (Valli *et al*., 2024*b*).

Correlation analyses for variables of interest were performed using Pearson correlation coefficient and p-value.

A feature importance analysis was conducted to identify which predictive variables (i.e., muscle fibre diameters and ion channel mRNA expression) had the most significant impact on the prediction of mean MU CV (the outcome variable). This analysis was performed using a random forest regressor model (Breiman, 2001) trained with leave-one-out cross-validation, which maximises the training data available at each iteration. In each iteration, the model was trained on all observations except one, optimizing parameters based on the remaining data. The importance of each predictive feature was assessed by its contribution to the model’s predictive accuracy. Results from all iterations were aggregated to calculate the mean importance score for each feature across all folds. This analysis estimates the relative significance of each predictive variable in explaining variability in mean MU CV. Feature importance analysis was conducted both including all data collection points (to estimate the importance of the predicting variables across the interventions) or only LS0 (to estimate the importance of the predicting variables at baseline).

The mixed model for MU CV was computed with jamovi 2.2.2 (Sydney, Australia – R language) while all the other statistical analyses were done using Python (V3.11.6, Python Software Foundation, USA, with the statsmodels v0.14.1 (Seabold & Perktold, 2010), scikit-learn v1.5.0 (Pedregosa *et al*., 2011) and pingouin v0.5.4 (Vallat, 2018) libraries). Statistical significance was accepted at p<0.05. The results are reported and plotted as mean (standard error) for linear models, and reported and plotted as mean (standard deviation) for the other analyses.

## 3. Results

### 3.1. Participants

Out of 12 participants, one dropped out after baseline measures for personal reasons. Eleven participants successfully completed the study without any adverse event. We were able to perform MU analysis on all the 11 participants, at all the data collection points. Instead, two participants opted to do not undergo muscle biopsies at AR21 and one participant was excluded from the muscle fibre diameters analysis at LS10 due to an insufficient amount of muscle sample.

### 3.2. Maximum voluntary contraction

For MVC, a main effect of time was observed (p<0.0001, □_p_^2^=0.8620). MVC decreased at LS10 (- 29.23%, p<0.0001, from 795.96 (112.83) N at LS0 to 563.37 (101.01) N at LS10) and returned to LS0 values at AR21 (p=0.834, 791.07 (107.06) N at AR21).

### 3.3. Motor unit decomposition and conduction velocity analysis

A total of 1542 unique MUs (654 at 10%, 577 at 25% and 311 at 50% MVC) were identified. Of these, 939 MUs (60.89% of the total pool, 332 at 10%, 383 at 25% and 224 at 50% MVC) were suitable for MU CV analysis. This resulted in an average number of MUs per participant and per data collection point of 10.37 (5.28) at 10%, 11.96 (6.00) at 25% and 7.47 (3.89) at 50% MVC.

On average, MU CV has been estimated on 4.00 (0.84) channels for each MU with a very high XCC between the selected channels of 0.95 (0.03) at LS0, 0.96 (0.02) at LS10 and 0.95 (0.03) at AR21.

MU CV (Fig. 2A) showed a significant Time effect (p<0.0001). MU CV was reduced at LS10 compared to LS0 (p<0.0001) while it increased at AR21 up to exceeding the LS0 values (p<0.0001). All the contraction intensities exhibited similar changes (no Time*Intensity effect, p=0.4103) although MU CV differed between contraction intensities (Intensity effect, p<0.0001). Estimated MU CV values and Post Hoc tests are presented in Table 1.

**Figure 2:**
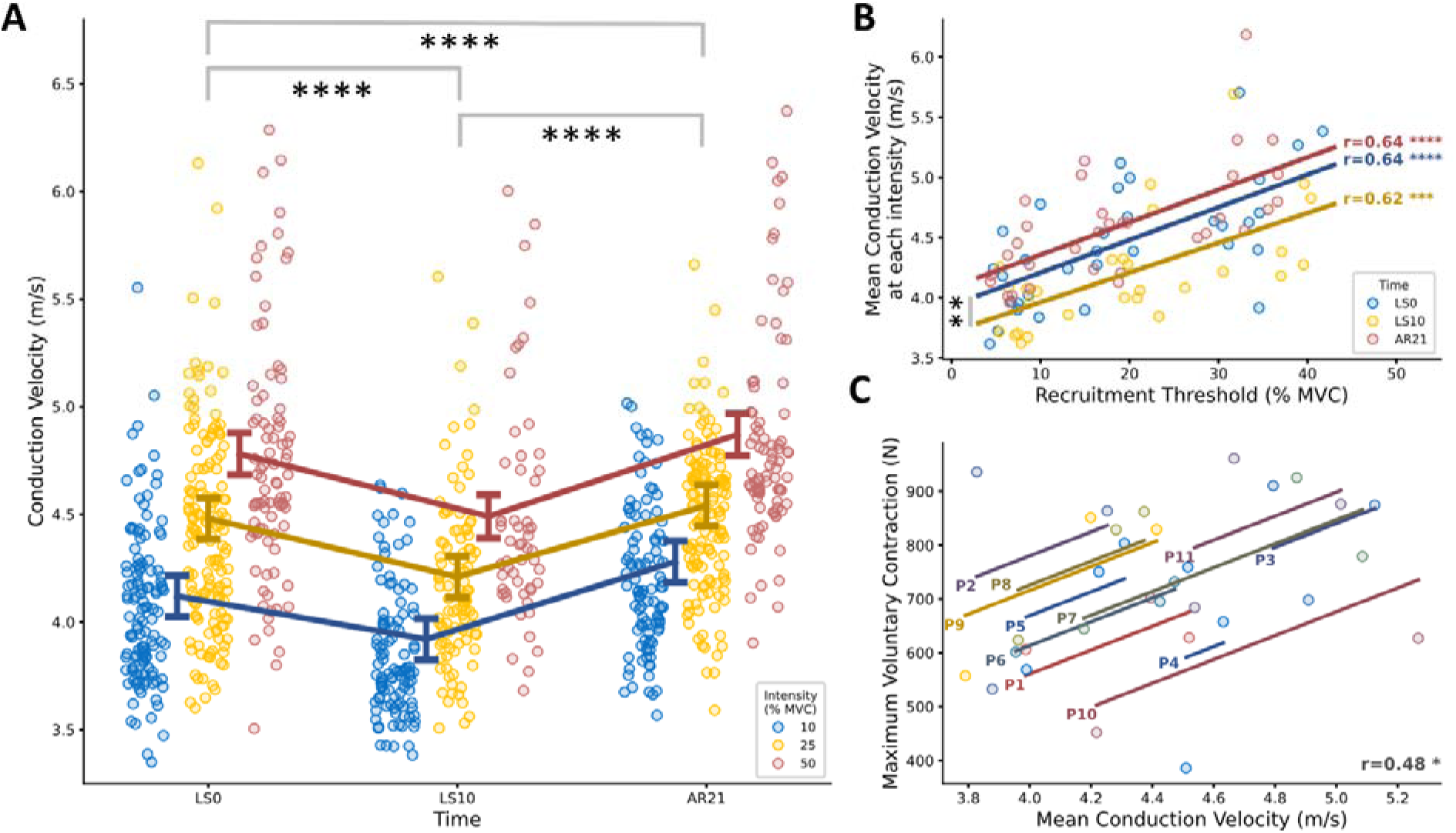
Analysis of motor unit conduction velocity (MU CV) and its association with MU recruitment threshold (RT) and maximum voluntary contraction (MVC). (A) Swarm plot representing MU CV at the three data collection points. Different intensities of contraction (i.e., 10, 25 and 50% MVC) are represented by different colours. Single MUs are represented by dots. Summary data are presented as mean ± standard error and the direction of the changes is highlighted by a connection line. Significant differences between the data collection points have been marked. (B) Correlation analysis between mean relative MU RT and CV. The correlation has been performed at each data collection point (represented by different colours) on mean MU RT and CV values. Mean values have been estimated for each participant at each contraction intensity (i.e., 3 values per participant, per data collection point). Significant differences between the intercept of the regression lines have been marked. (C) Common within-individual association between MU CV and MVC across the data collection points. Participants are represented by different colours. MVC, maximum voluntary contraction; LS0, day 0 of limb suspension; LS10, after 10 days of ULLS; AR21, after 21 days of active recovery. Significance levels are: *p<0.05, **p<0.01, ***p<0.001, ****p<0.0001.

**Table 1:**
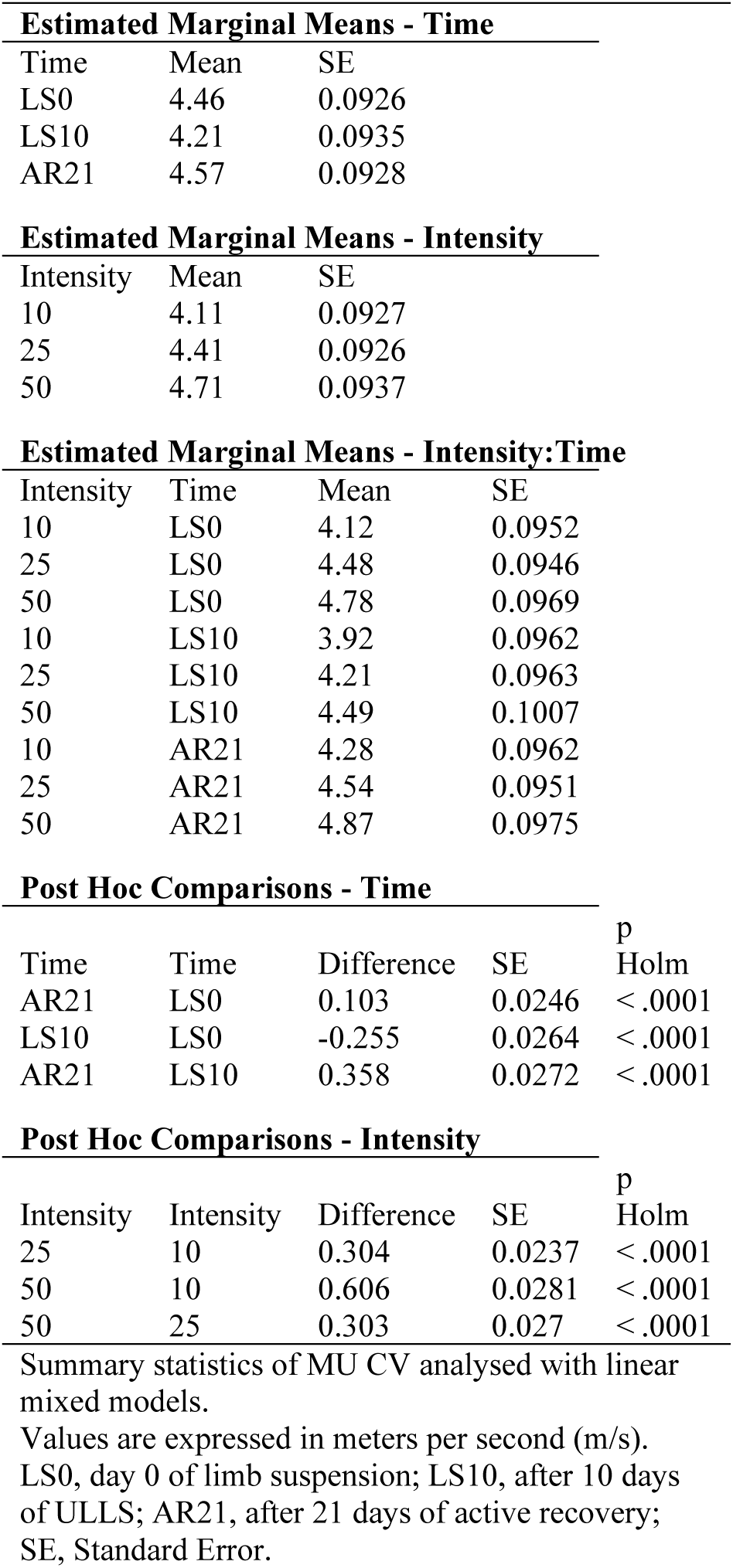
Motor unit conduction velocity (MU CV) Estimated Means and Post Hoc tests for significant effects.

| Estimated Marginal Means - Time |  |  |  |  |
| --- | --- | --- | --- | --- |
| Time | Mean | SE |  |  |
| LS0 | 4.46 | 0.0926 |  |  |
| LS10 | 4.21 | 0.0935 |  |  |
| AR21 | 4.57 | 0.0928 |  |  |
| Estimated Marginal Means - Intensity |  |  |  |  |
| Intensity | Mean | SE |  |  |
| 10 | 4.11 | 0.0927 |  |  |
| 25 | 4.41 | 0.0926 |  |  |
| 50 | 4.71 | 0.0937 |  |  |
| Estimated Marginal Means - Intensity:Time |  |  |  |  |
| Intensity | Time | Mean | SE |  |
| 10 | LS0 | 4.12 | 0.0952 |  |
| 25 | LS0 | 4.48 | 0.0946 |  |
| 50 | LS0 | 4.78 | 0.0969 |  |
| 10 | LS10 | 3.92 | 0.0962 |  |
| 25 | LS10 | 4.21 | 0.0963 |  |
| 50 | LS10 | 4.49 | 0.1007 |  |
| 10 | AR21 | 4.28 | 0.0962 |  |
| 25 | AR21 | 4.54 | 0.0951 |  |
| 50 | AR21 | 4.87 | 0.0975 |  |
| Post Hoc Comparisons - Time |  |  |  |  |
| Time | Time | Difference | SE | p Holm |
| AR21 | LS0 | 0.103 | 0.0246 | <□.0001 |
| LS10 | LS0 | -0.255 | 0.0264 | <□.0001 |
| AR21 | LS10 | 0.358 | 0.0272 | <□.0001 |
| Post Hoc Comparisons - Intensity |  |  |  |  |
| Intensity | Intensity | Difference | SE | p Holm |
| 25 | 10 | 0.304 | 0.0237 | <□.0001 |
| 50 | 10 | 0.606 | 0.0281 | <□.0001 |
| 50 | 25 | 0.303 | 0.027 | <□.0001 |

Summary statistics of MU CV analysed with linear mixed models.
Values are expressed in meters per second (m/s).
LS0, day 0 of limb suspension; LS10, after 10 days of ULLS; AR21, after 21 days of active recovery; SE, Standard Error.

MU RT and CV were significantly correlated at each data collection point (r=0.64, p<0.0001 at LS0, r=0.62, p=0.0004 at LS10 and r=0.64, p<0.0001 at AR21) (Fig 2B). No significant differences have been detected between the slopes of the regression lines (p=0.7714 for LS0 vs LS10 and p=0.9882 for LS0 vs AR21). The intercepts of the regression lines differed between LS0 and LS10 (p=0.0059) but not between LS0 and AR21 (p=0.1136).

Changes across data collection points in MU CV significantly correlated with changes in MVC (r=0.48, p=0.0236) (Fig 2C).

### 3.4. Muscle fibre diameters

Muscle fibre diameters did not change across the intervention, regardless of fibre type. Slow fibres had a diameter of 51.43 (6.95) µm at LS0, 49.81 (10.49) µm at LS10 and 47.43 (7.10) µm at AR21 (p=0.3102, □_p_^2^=0.1540) (Fig. 3A). Fast fibres had a diameter of 54.17 (6.50) µm at LS0, 52.46 (9.01) µm at LS10 and 51.81 (7.90) µm at AR21 (p= 0.7106, □_p_^2^=0.0476) (Fig. 3B). In total, fibres had a diameter of 53.30 (6.30) µm at LS0, 51.60 (8.16) µm at LS10 and 49.61 (7.60) µm at AR21 (p= 0.3953, □_p_^2^=0.1242) (Fig. 3C).

**Figure 3:**
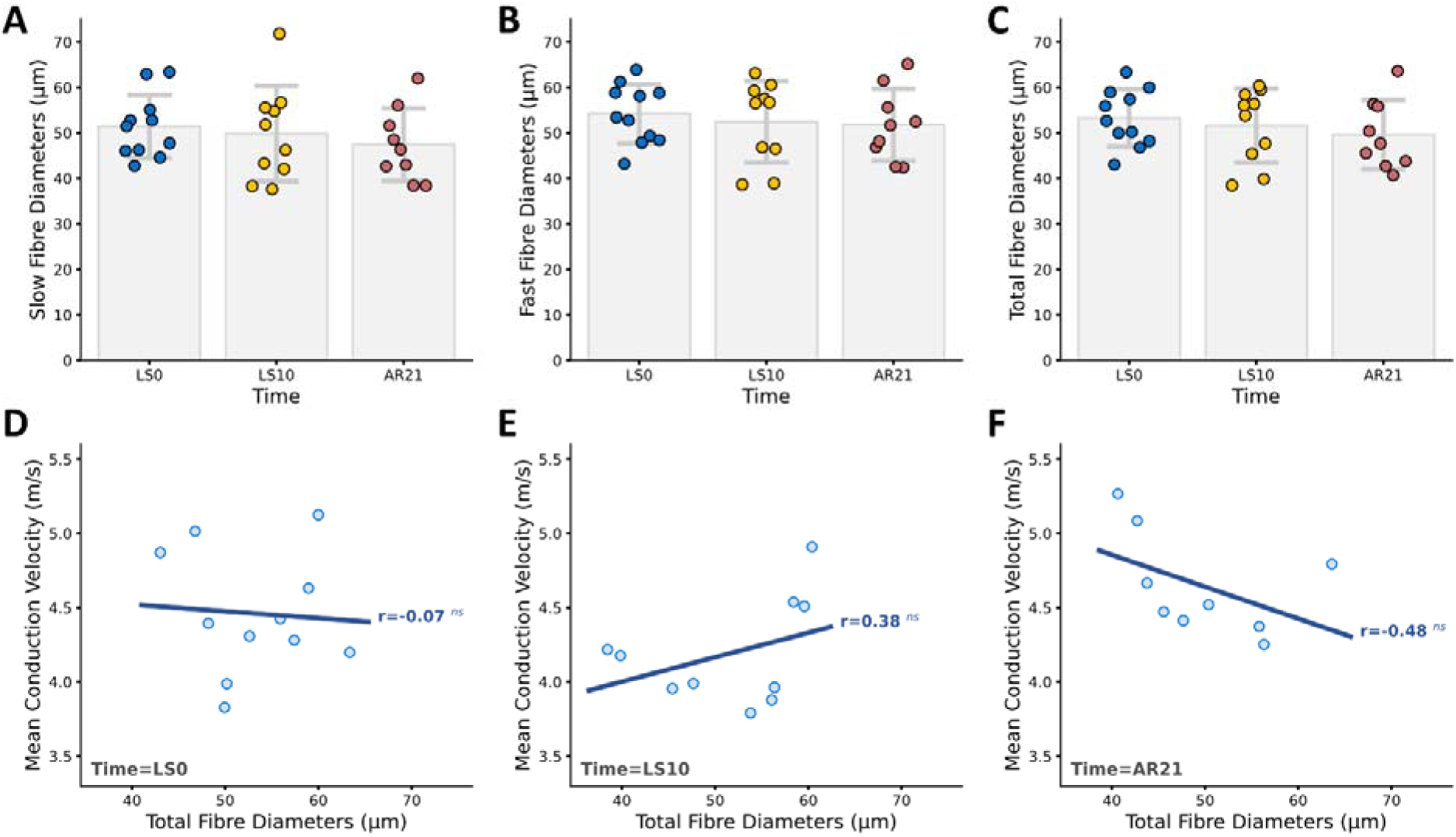
Analysis of muscle fibre diameters and their association with motor unit conduction velocity (MU CV). Scatter plot and box plot representing individual values and mean ± standard deviation of slow (A), fast (B) and total fibre diameters at each data collection point. Scatter plot showing the correlation between total fibre diameters and mean MU CV at each data collection point. LS0, day 0 of limb suspension; LS10, after 10 days of ULLS; AR21, after 21 days of active recovery. Significance levels are: ^ns^p>0.05.

There was no significant correlation between total fibre diameters and mean MU CV values at each data collection point, as demonstrated by the correlation analyses (r=-0.07, p=0.8367 at LS0, r=0.38, p=0.2812 at LS10, r=-0.48, p=0.1928 at AR21) (Fig. 3D-F).

Similarly, there was no significant correlation between changes in total fibre diameters and mean MU CV values across the data collection points, as demonstrated by repeated measures correlation analysis (r=0.01, p=0.9675) (not shown in figure).

### 3.5. Ion channels mRNA expression

The feature importance analysis conducted including ion channel mRNA expression and muscle fibre diameters at all data collection points revealed that the top four features (i.e., the ion channels KCNJ2-AS1, KCNN2, KCNN3 and SCN4A) accounted for 49% of the predictive importance for MU CV. Notably, potassium channels exhibited a dominant influence, with the first three channels together contributing to 40% of the overall importance. A comprehensive overview of all features and their respective importance scores is presented in Fig. 4A.

**Figure 4:**
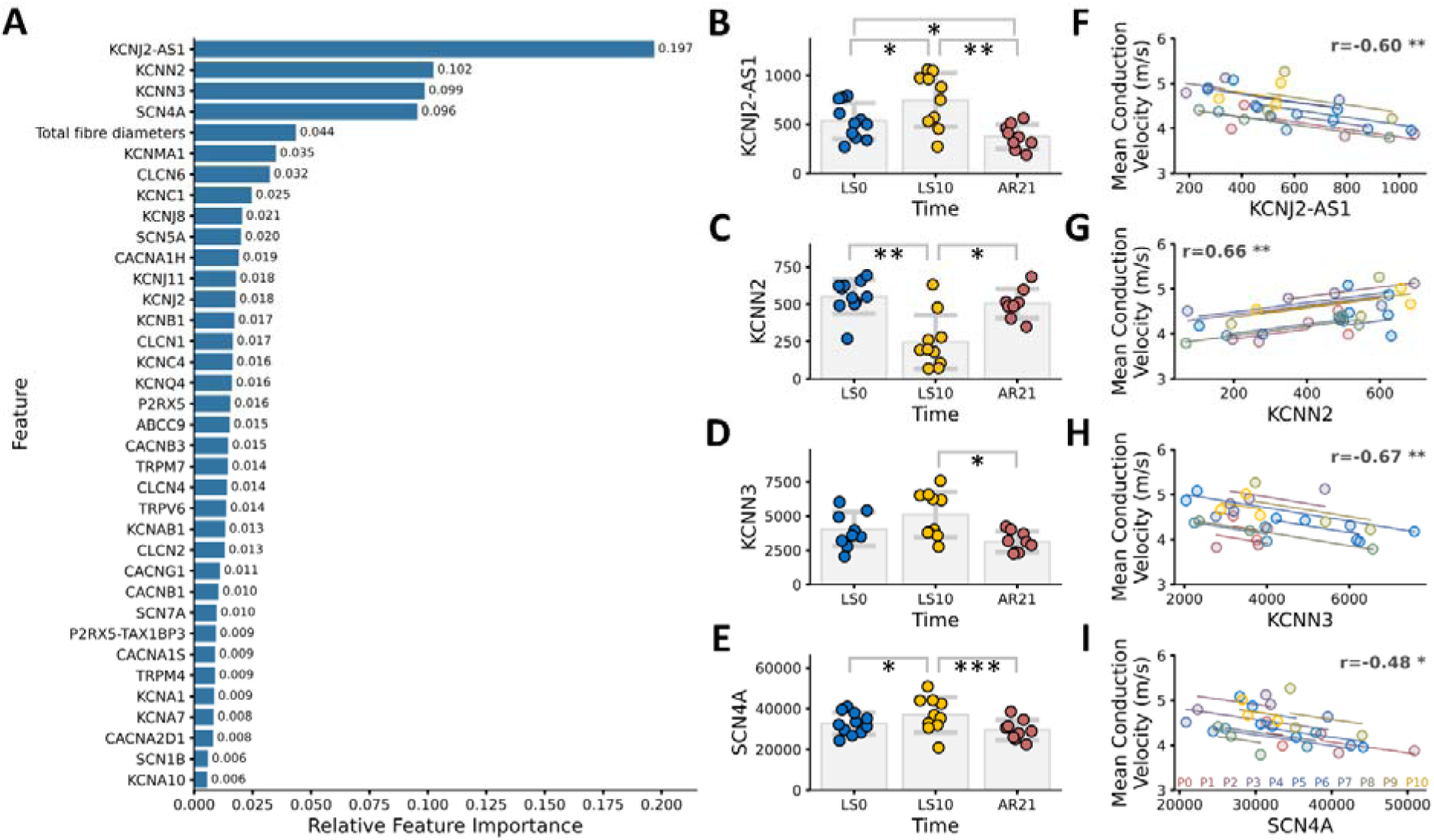
Analysis of ion channel mRNA expression levels and their association with motor unit conduction velocity (MU CV) across the data collection points. Bar plot representing the relative importance of each feature (ion channel genes and total muscle fibre diameters) for the prediction of motor unit conduction velocity (MU CV) (A). Scatter plot and box plot representing individual values and mean ± standard deviation of KCNJ2-AS1 (B), KCNN2 (C), KCNN3 (D) and SCN4A (E) mRNA expression levels (normalised reads count) at each data collection point. Common within-individual association between KCNJ2-AS1 (F), KCNN2 (G), KCNN3 (H) and SCN4A (I) mRNA expression levels and mean MU CV across the data collection points. Participants are represented by different colours. LS0, day 0 of limb suspension; LS10, after 10 days of ULLS; AR21, after 21 days of active recovery. Significance levels are: *p<0.05, **p<0.01, ***p<0.001.

The changes in the mRNA expression levels (normalised reads count) of the top four ion channel genes across data collection points has also been investigated. For the mRNA expression level of KCNJ2-AS1 (Fig. 4B), a main effect of time was observed (p=0.0003, □_p_^2^=0.6927). KCNJ2-AS1 increased at LS10 (from 533.37 (183.72) at LS0 to 748.98 (277.31) at LS10, p=0.0401) and decreased at AR21 (to 373.27 (124.45), p=0.0032 for LS10 vs AR21 and p=0.0401 for LS0 vs AR21). For the mRNA expression level of KCNN2 (Fig. 4C), a main effect of time was observed (p=0.0016, □_p_^2^=0.6014). KCNN2 decreased at LS10 (from 551.70 (115.16) at LS0 to 246.54 (180.34) at LS10, p=0.0090) and returned to baseline at AR21 (to 503.66 (97.62), p=0.0462 for LS10 vs AR21). For the mRNA expression level of KCNN3 (Fig. 4D), a main effect of time was observed (p=0.0137, □_p_^2^=0.4581). KCNN3 decreased at AR21 (from 5106.52 (1670.60) at LS10 to 3115.98 (746.13) at AR21, p=0.0414). For the mRNA expression level of SCN4A (Fig. 4E), a main effect of time was observed (p=0.0001, □_p_^2^=0.7376). SCN4A increased at LS10 (from 32691.57 (5322.64) at LS0 to 37000.38 (8630.74) at LS10, p=0.0208) and returned to baseline at AR21 (to 29512.52 (4981.95), p=0.0003 for LS10 vs AR21).

Changes in the mRNA expression levels of all four top-ranked ion channel genes showed significant correlations with changes in mean MU CV throughout the intervention, as demonstrated by repeated measures correlation analyses (r=-0.60, p= 0.0048 for KCNJ2-AS1, r=0.66, p=0.0014 for KCNN2, r=-0.67, p=0.0011 for KCNN3, r=-0.48, p=0.0331 for SCN4A) (Fig. 4F-I).

The feature importance analysis conducted including only ion channel mRNA expression at LS0 revealed that the top four ion channels (KCNN2, KCNC1 KCNA7 and KCNN3, ordered for importance) accounted for 51% of the predictive importance for MU CV. An overview of all features and their respective importance scores is presented in Fig. 5A.

**Figure 5:**
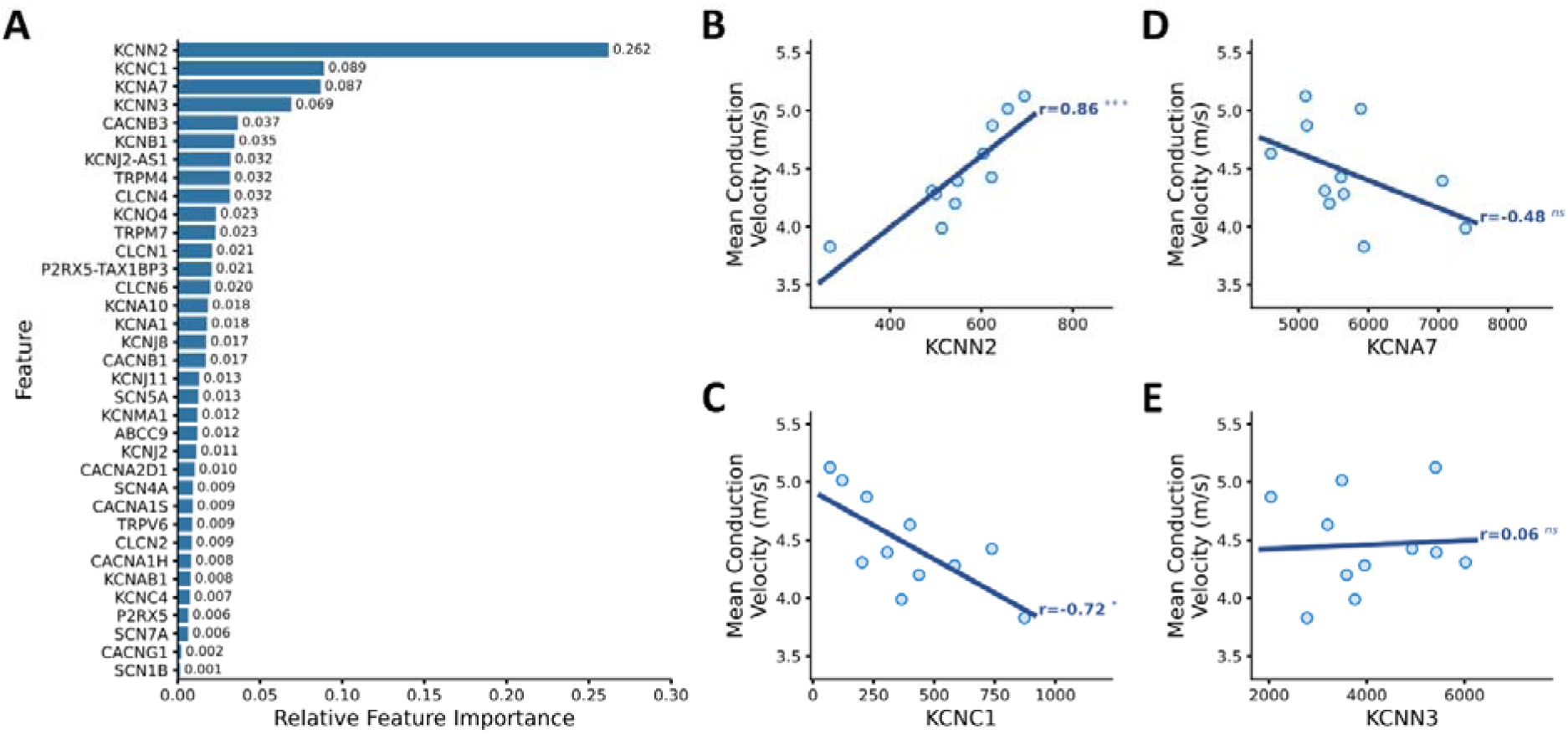
Analysis of ion channel mRNA expression levels and their association with motor unit conduction velocity (MU CV) at day 0 of limb suspension (LS0, baseline). Bar plot representing the relative importance of each feature (ion channel genes) for the prediction of motor unit conduction velocity (MU CV) (A). Scatter plot showing the correlation between KCNN2 (B), KCNC1 (C), KCNA7 (D) and KCNN3 (E) mRNA expression levels (normalised reads count) and mean MU CV. Significance levels are: ^ns^p>0.05, *p<0.05, ***p<0.001.

KCNN2 and KCNC1 mRNA expression levels correlated with mean MU CV at LS0 (r= 0.86, p= 0.0007, for KCNN2, r=0.72, p=0.0129 for KCNC1) while KCNA7 and KCNN3 did not show significant correlations (Fig. 5B-E).

## 4. Discussion

This study aimed to investigate the role of changes in motor unit conduction velocity (MU CV) in muscle dysfunction following disuse and active recovery, as well as the potential biological factors underlying these changes. MU CV was significantly reduced following 10 days of unloading of the dominant lower limb (ULLS) in young, healthy males. Remarkably, MU CV fully recovered, and even exceeded baseline levels, following 21 days of active recovery based on resistance exercise. The observed changes in MU CV correlated with alterations in maximal force (MVC), suggesting that the reduction in MU CV may be a determinant of muscle function loss during periods of unloading or disuse. While no correlation was observed between MU CV and muscle fibre diameters, mRNA expression levels of a small subset of skeletal muscle ion channel genes, particularly those linked to K^+^ transport, could predict ∼50% of the variability in MU CV at baseline and throughout the intervention. Although the role of ion channels in action potential generation and propagation is well established, this study is the first to directly relate specific ion channel mRNA expression levels to MU CV in humans, suggesting that ion channels may play a pivotal role in determining MU CV at baseline and it’s changes throughout the interventions, beyond the influence of muscle fibre size.

### 4.1. Effect of the interventions on motor unit conduction velocity

ULLS induced a decline in MU CV, with values at LS10 significantly lower than those at LS0 (Fig. 2A). This finding is consistent with previous observations by (Cescon & Gazzoni, 2010), who reported similar results after 14 days of bed rest. Notably, this decline was consistent across all contraction intensities, indicating that the impairment of MU CV affects all MUs uniformly. This finding suggests a generalized decline in the muscle unit function following 10 days of ULLS impacting both slower and faster MUs equally. These observations align with our previous hypothesis that short-term disuse would impair the function of all muscle units, regardless of their RT while selectively reducing the discharge rate of lower-threshold MUs (Valli *et al*., 2024*b*). This was initially supported by literature showing that short periods of bed rest and ULLS impair muscle fibre contractility by altering intracellular calcium handling (Monti *et al*., 2021) and reducing fibre-specific force (Brocca *et al*., 2015) in both slow and fast muscle fibres. This study further confirms this uniform impairment given that similar changes in MU CV are observed across all contraction intensities. Overall, these findings offer additional support to our previous hypothesis that changes in the neural drive to muscle likely precede the metabolic adaptations associated with muscle fibre-type shifts (Valli *et al*., 2024*b*), such as the transition from slow to fast fibres, which are commonly observed after prolonged periods of muscle disuse (Hortobágyi *et al*., 2000; Ciciliot *et al*., 2013).

The disuse-induced reduction in MU CV observed at LS10 was fully reversed by AR21, with MU CV surpassing the baseline levels (Fig. 2A). Since MU CV changed across the intervention period, we examined whether the interventions could also induce changes in the relationship between MU RT and CV (Arendt-Nielsen *et al*., 1984). At LS0, MU CV values positively correlated with RT, where higher RT values corresponded to higher MU CV. This relationship remained stable over the intervention period, as evidenced by the similar slopes of the regression lines at each time point (Fig. 2B), suggesting that the underlying physiological mechanisms linking MU CV and RT were unaffected by ULLS or AR.

Lastly, we observed that changes in MU CV correlated with the changes in MVC (Fig. 2C), suggesting that reductions or increases in MU CV are linked to corresponding declines or improvements in muscle force. This relationship underscores the potential importance of MU CV as an indicator of muscle function and highlights that, understanding the mechanisms governing MU CV, could provide novel insights on the mechanisms responsible for the loss and recovery of muscle function after muscle disuse/unloading and recovery. In light of this, we investigated whether changes in structural or molecular factors such as muscle fibre diameters and skeletal muscle ion channel mRNA expression could be related to changes in MU CV across the intervention.

### 4.2. Muscle fibre diameters and their relationship with motor unit conduction velocity

Muscle fibre diameters, whether categorized as slow, fast, or total, showed no significant change across any data collection points (Fig. 3A-C). This stability in muscle fibre size following short-term disuse is consistent with previous findings from immobilisation and bed rest studies, which often report limited changes in muscle fibre size (Hespel *et al*., 2001; Pišot *et al*., 2016; Reidy *et al*., 2018; Monti *et al*., 2021). However, when disuse models involve more severe conditions, such as cast immobilisation or dry immersion, muscle fibre atrophy may occur even within short timeframes (i.e., 3-7 days) (Suetta *et al*., 2012; Wall *et al*., 2014; Demangel *et al*., 2017). It should be noted that longer periods of disuse consistently result in a reduction of muscle fibre size (Campbell *et al*., 2013; Brocca *et al*., 2015), emphasizing that muscle fibre size changes depend on both the severity and duration of the disuse model. Therefore, given the relatively short 10-day period of ULLS in the current study, the lack of change in muscle fibre diameters that we observed is consistent with the literature.

With the aim of identifying the factors regulating MU CV, we tested whether MU CV was correlated with muscle fibre diameters, both at baseline and across data collection points. Indeed, previous studies observed a clear correlation between CV of electrically evoked action potentials and muscle fibre size in humans (Blijham *et al*., 2006; Methenitis *et al*., 2016). Interestingly, we found no correlation between MU CV and muscle fibre diameters, both within and across data collection points.

We hypothesise that the absence of correlation observed at baseline, is likely due to the type of investigated action potentials, given that we studied the CV of action potentials generated by voluntarily activated MUs while Blijham and Methenitis studied the CV of electrically evoked action potentials. As a matter of fact, CV values vary depending on whether action potentials are generated via electrical stimulation or voluntary contraction, with the latter typically being faster and more variable (Buchthal *et al*., 1955). Furthermore, during electrical stimulation of the muscle, only fibres located close to the electrode are activated. These fibres tend to have nearly identical membrane potentials, resulting in a highly uniform biophysical response (Buchthal *et al*., 1955). Indeed, the stimulated fibres are activated because of spatial and electrical properties, without any guarantee of belonging to the same MU (Arendt-Nielsen & Zwarts, 1989).

Therefore, while muscle fibre diameter is probably an important factor in determining muscle fibre CV, the relationship appears more complex, particularly in the context of voluntarily activated MUs, which involve multiple, spatially distributed fibres and may not directly reflect the properties observed under electrical stimulation. To our knowledge, the only study reporting a direct comparison between voluntary muscle fibre CV and muscle fibre diameters aligns with our findings, as it found no significant correlation between the two parameters in the *vastus lateralis* muscle (Sadoyama *et al*., 1988).

Across the interventions, it could be expected that changes in MU CV do not relate to changes in muscle fibre diameters. Indeed, even acute metabolic modifications such as the increased buffering capacity provoked by sodium bicarbonate ingestion (Hunter *et al*., 2009) or muscle fatigue (Vila-Chã *et al*., 2012) can induce substantial alterations in CV without affecting muscle fibre diameters. Similarly, just two weeks of high-intensity interval training or endurance exercise can cause substantial MU CV changes, yet such a short intervention is unlikely to induce measurable changes in fibre diameters (Martinez-Valdes *et al*., 2018).

In conclusion, the findings of our study together with the existing literature show that MU CV and its changes, at least in the *vastus lateralis* muscle, do not correlate with muscle fibre size. This finding implies the presence of other factors with a significant role in determining MU CV and its changes.

### 4.3. Ion channels and their relationship with motor unit conduction velocity

The generation and propagation of action potentials in muscle fibres are regulated by ion flux across the membrane (Jurkat-Rott & Lehmann-Horn, 2004). Indeed, the depolarisation phase is initiated by the opening of voltage-gated sodium channels, allowing sodium ions (Na^+^) to flow into the cell, leading to a rapid change in the membrane’s electrical charge. The subsequent repolarisation phase is driven by the opening of potassium channels, which facilitate the efflux of potassium ions (K^+^), restoring the membrane’s negative charge. Eventually, after the conclusion of the action potential, the Na^+^/K^+^ ATPase pump restores ion gradients by pumping Na^+^ out and K^+^ into the cell, preparing the muscle fibre for the following action potential (Feher, 2017). Maintaining an optimal balance of Na^+^ and K^+^ ions is essential for muscle excitability and contractility (Nielsen & Clausen, 2000) and the involved structures can undergo both structural and functional adaptations in response to different physical demands (Green *et al*., 2004).

Given the role of skeletal muscle ion channels in the generation and propagation of action potentials, we conducted a feature importance analysis to assess the predictive significance of the mRNA expression levels of 35 ion channels in explaining MU CV across the interventions (Fig. 4A). Muscle fibre diameters were also included in this analysis as a reference in order to compare their contribution. The results of the feature importance analysis highlighted four key ion channel genes (*KCNJ2-AS1*, *KCNN2*, *KCNN3* and *SCN4A*) as the most influential for the prediction of MU CV, accounting for 49% of the model’s predictive importance. Notably, the first three genes, responsible for 40% of the importance, encode potassium channels, while *SCN4A* encodes a sodium channel. In contrast, muscle fibre diameters contributed marginally, with only 4.4% of the predictive importance. Interestingly, the mRNA expression level of these key ion channel genes was modulated by the interventions (Fig. 4B-E) and correlated significantly with MU CV values (Fig. 4F-I). These novel findings highlight the importance of potassium and sodium channels in the modulation of MU CV, and suggest that ion channels may have a far more critical role in regulating MU CV than muscle fibre size, at least in an anatomically complex muscle such as the *vastus lateralis*.

The top-ranked ion channel gene, *KCNJ2-AS1*, encodes an antisense RNA of *KCNJ2*. This antisense RNA may play a regulatory role in the expression or activity of KCNJ2, which encodes the Kir2.1 inwardly rectifying K^+^ channel (Hibino *et al*., 2010). Kir2.1 contributes to the establishment of highly negative membrane potential and long-lasting hyperpolarization, crucial for rapid Na^+^ channel activation during depolarisation, which is central to action potential initiation and propagation (Hibino *et al*., 2010).

Alterations in Kir2.1 function are implicated in Andersen’s syndrome, where dysregulation of K^+^ rectification disrupts Na^+^ channel dynamics, impairing both action potential generation and propagation, eventually resulting in periodic paralysis (Plaster *et al*., 2001). Furthermore, Kir2.1 has a key role in the regulation of the differentiation and fusion of myoblasts to form a multinucleated muscle fibre, with mutations in the gene resulting in muscle weakness (Fischer-Lougheed *et al*., 2001; Konig *et al*., 2004). Similar consequences are observed also for mutations in the sodium channel Na_v_1.4, which is encoded by the *SCN4A* gene (Ptácek *et al*., 1991).

To understand whether ion channel mRNA expression correlates with MU CV also in physiological conditions at baseline, we repeated the feature importance analysis including only ion channel genes at LS0 (Fig. 5A). Indeed, transcriptional changes observed after interventions may not directly reflect the pathways modulating CV in a resting, unaltered state. Interestingly, although the specific gene rankings differed when considering only LS0, potassium channel genes continued to dominate in importance. The top four genes (*KCNN2*, *KCNC1*, *KCNA7*, and *KCNN3*) accounted for 51% of the predictive power for MU CV at LS0. Among these, *KCNN2* showed a particularly strong correlation with MU CV values (r = 0.86, Fig. 5B), suggesting that this gene plays a crucial role in MU CV regulation both at rest and during the intervention phases.

The *KCNN2* gene belongs to the KCNN family, which includes *KCNN1*, *KCNN2*, and *KCNN3*, encoding the small conductance calcium-activated potassium (SK) channels KCa2.1, KCa2.2 and KCa2.3, respectively (Köhler *et al*., 1996). While KCa2.1 is usually not expressed in muscles (Chen *et al*., 2004), KCa2.2 and KCa2.3 channels are activated by increases in intracellular Ca^2+^ and regulate the afterhyperpolarisation phase of the action potential, both in neurons and muscle fibres (Guéguinou *et al*., 2014; Rahman *et al*., 2023). Although these ion channels are little studied in humans, it has been observed that in the rat soleus muscle both the KCa2.3 protein and the *KCNN3* mRNA expression are upregulated upon denervation (Favero *et al*., 2008), Interestingly, early indications of denervation were observed also in our cohort at LS10 (Sarto *et al*., 2022*a*), with a concomitant increase in the *KCNN3* mRNA expression (Fig. 4D).

In neurons, changes in the duration of the afterhyperpolarisation phase have been correlated with changes in CV, with longer afterhyperpolarisation periods usually associated with lower CV values (Gardiner & Kernell, 1990; Cross & Robertson, 2016). These findings align well with simulation studies in human muscles showing that muscle fibre CV can be modulated by altering the afterhyperpolarisation phase through changes in KCa channels activity (Fortune & Lowery, 2011). Given that the Ca^2+^ activating the KCa channels in skeletal muscle is released from the sarcoplasmic reticulum (Neelands *et al*., 2001), alterations in Ca^2+^ release could affect the activity of KCa channels and, consequently, MU CV. Evidence has shown that 10 days of bed rest reduces the Ca^2+^ content in the sarcoplasmic reticulum and diminishes the responsiveness of calcium release channels (Monti *et al*., 2021). This suggests that reductions in Ca^2+^ availability, due to sarcoplasmic reticulum dysfunction, could impair KCa channel function and modulate MU CV during periods of muscle disuse.

In summary, our findings suggest that Kir2.1 may modulate MU CV by influencing the generation of action potentials, while KCa channels affect the afterhyperpolarisation phase. Furthermore, the functioning of KCa channels suggests a dynamic relationship between calcium-handling mechanisms, KCa channel activity, and MU CV. The modulation of the afterhyperpolarisation phase by KCa channels in response to intracellular Ca^+^ changes offers novel insights into how ion channels may regulate MU CV beyond structural determinants.

### 4.4. Methodological considerations

Limitations in study design, HDsEMG recordings and muscle sampling have already been discussed in (Sarto *et al*., 2022*a*; Valli *et al*., 2024*b*).

To ensure the most accurate estimation of MU CV in this study, several precautions have been used.

First, the electrode grid was carefully aligned along the muscle fibre direction, identified objectively through B-mode ultrasound imaging. Second, the estimation of MU CV was performed using state-of-the-art algorithms (Farina *et al*., 2002; Farina & Merletti, 2004) and channel selection was executed with rigorous quality criteria, as indicated by the high XCC values obtained (Valli *et al*., 2024*a*). However, the architecture of the *vastus lateralis* muscle, with its pennate structure and multi-angled fibre arrangement, may introduce some variability in MU CV estimation compared to fusiform muscles like the *biceps brachii* (Casolo *et al*., 2023). Therefore, this anatomical variability should be considered when comparing MU CV findings across studies that focus on different muscle groups, as these structural differences may influence MU CV estimates.

While mRNA expression analysis provides valuable insights into rapidly changing molecular processes, it may not fully reflect the quantity and activity of the corresponding proteins. This discrepancy arises due to the complex regulatory mechanisms that occur post-transcriptionally, which include protein translation, degradation, and post-translational modifications, all of which determine the final protein content and functionality. Indeed, proteins often undergo modifications (e.g., phosphorylation, glycosylation) that can alter their activity without changes in mRNA expression. These modifications are essential in functional regulation, particularly in ion channels, where changes in phosphorylation state can impact channel opening, closing, and sensitivity to ions (Jurkat-Rott & Lehmann-Horn, 2004). Additionally, the localisation of these proteins within muscle cells can influence their accessibility and function during muscle activation, as seen in various ion channels whose distribution affects muscle fibre excitability (Tricarico *et al*., 1997). Therefore, to reinforce and deepen the findings of this study, future investigations should focus on quantifying protein content and activity levels, assessing post-translational modifications, and determining spatial distribution within muscle fibres.

Ultimately, although our study identified associations between certain ion channel genes and MU CV, much remains unknown about the specific roles these genes play in skeletal muscle and their practical influence on MU CV modulation. Therefore, further research focusing on the individual contributions of these ion channels in muscle tissue is essential to fully contextualise our findings and clarify their impact on MU CV.

## Abbreviations

AR: Active recovery
AR21: Day 21 of active recovery
CV: Conduction Velocity
EMG: Electromyography
HDsEMG: High-Density surface Electromyography
LS0: Day 0 of limb suspension
LS10: day 10 of limb suspension
MU: Motor Unit
MUAP: Motor Unit Action Potential
MVC: Maximum Voluntary Contraction
RT: Recruitment Threshold
SK: Small conductance calcium-activated Potassium channels
ULLS: Unilateral Lower Limb Suspension
XCC: Cross-Correlation Coefficient

